# Expansions of Price Equation for Viability and Fecundity Selection

**DOI:** 10.1101/2024.06.24.600333

**Authors:** Jia-Xu Han, Rui-Wu Wang

## Abstract

The Price equation offers a means of separating the portion of population change attributable to natural selection from the overall change. There is debate over the universality of the Price equation due to the belief that it makes simple assumptions. However, it should be noted that the definition of fitness is assumed within the Price equation. To account for populations subject to viability selection and fecundity selection during their life cycle, two expansions of the Price equation have been proposed based on a clear understanding of fitness. These expansions include three terms: a viability selection term, a fecundity selection term, and a mean value term that captures the difference between adults and their zygotes. Unlike the classic Price equation, which also has a mean value term capturing the difference between adults and their successful zygotes, the mean value term in our expansions will be zero in the absence of mutation, drift, and recombination. Furthermore, the individual-based simulation shows that our expansions can outperform the classic Price equation in capturing the average change in strategy for resource allocation. In summary, our expansions fully capture the effects of natural selection and separate them into viability and fecundity selection terms. Our expansions allows for a wider application of the Price equation, especially in the case where there is a trade-off between viability and fecundity selection.

## 1 Introduction

A crucial query in the field of evolutionary biology concerns the extent to which natural selection drives evolutionary responses. To address this question, it is necessary to differentiate the role of natural selection from variations in measurements such as fitness, gene frequency, and traits. A common method is to utilize a regression model that characterizes individual predictors and their contributions (Fisher 1918, Fisher 1958). Fisher’s Fundamental Theorem of Natural Selection is a particularly significant approach in this regard:

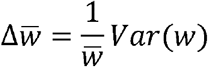

where w is fitness of individuals in a population. The Fundamental Theorem of Natural Selection asserts that the rate of increase of mean fitness in a population, as defined by a regression model, is exactly equal to the additive genetic variance in population fitness at any given moment (Fisher 1930; Edwards 2014). Despite the formulation of the theorem, there has been much debate over its interpretation and generalizability, with various scholars offering different perspectives on the matter by considering changes in mean genotypic fitness (Kempthorne 1957), changes in environment and genetic interactions (Kimura 1958), change in gene frequency (Price 1972), and the additive estimate part in fitness (Ewens 1989; Edwards 1994; Lessard 1997). One of the most significant works in this area is the Price equation, which rigorously examines Fisher’s statement as a theorem (Price 1970).

By defining fitness as a successful gametes/offspring, the Price equation express as

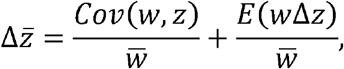

where *z* represents any trait of individuals, *w* represents fitness of individuals, Δ*z* represents the difference of the trait between the parent and its offspring and overbar denote population average. The first term in the Price equation reflects the fundamental concept of “survival of the fittest” proposed by Darwin, since it is positive only when there is a positive correlation between the trait value and the fitness. In other words, if we disregard the difference between an ancestor and the average of all its descendants, increasing the value of a trait that can enhance fitness or the number of successful gametes (or offspring) will result in an increase in the population’s mean trait value, and vice versa (Frank, 2012). The second term describes changes in trait values between parents and their offspring caused by various processes such as mutation and drift. When the trait is also fitness and there is no random drift, Fisher’s theorem is a special case of the Price equation.

The Price equation is widely used because it makes simple assumptions, which contributes to its popularity. However, this characteristic has also led to some debate. One perspective argues that because the Price equation is free from assumptions, it is always accurate (Frank 1997, 2012; Grafen 2002, 2020; Gardner 2020). Additionally, the Price equation has found extensive applications in theoretical biology, including the derivation of numerous fundamental theorems in population genetics (Grafen 2002, Queller 2017) and serving as the formal basis for both kin selection and group selection theories (Frank 1995; Gardner 2008). Conversely, some believe that both sides of the equation represent the same thing, rendering the equation meaningless and a mere mathematical tautology. This is because the absence of additional assumptions in the Price equation results in the right side of the equation being a redundant restatement of the left side (Nowak & Highfield 2011; van Veelen et al. 2012; van Veelen 2020).

To date, there have been few expansions to the Price equation, with some notable exceptions including those addressing fluctuating environments (Grafen 1999, 2000), finite populations (Rice 2008), and general mappings between ancestral and descendant populations (Kerr & Godfrey-Smith 2009). In this study, we propose a new expansion of the Price equation by redefining fitness and breaking down the selection component of the equation into viability selection and fertility selection. We begin by exploring two different measures of fitness before presenting our expansion to the equation, which is based on a more precise definition of fitness. Then, we compare the efficiency of the classic and expanded Price equations using individual-based simulations.

## 2 The definition of fitness

The presence of differences in fitness is a necessary condition for natural selection to occur and for organisms to adapt to their environment. Fitness is a metric used to evaluate the capacity of individuals, populations, or species to survive and reproduce in their specific environment (Orr 2009). As a result of their survival and reproduction, organisms transmit their genes or traits to the next generation. A common mathematical measure of absolute fitness is the expected number of copies of genes or traits left after a life cycle.

Consider a species with a basic life cycle. Zygotes are produced, and they either survive to adulthood or do not. If they do, adults try to court and mate. If successful, these adults generate a specific number of offspring, and the cycle restarts. Variations in fitness among individuals can emerge from disparities in performance at any of these stages. Each of these fitness elements, such as viability, fecundity, and mating success, can contribute to differences in overall fitness among individuals, which can cause different individuals to leave different numbers of offspring. Here, we only consider the role of natural selection and disregard sexual selection, which implies that the effect of mating success is overlooked. Then, in each life cycle, there is a selection for viability of zygotes and a selection for fecundity of adults.

When considering the ancestral and descendant populations in the Price equation, there are two fitness measurements depending on the order of viability selection and fecundity selection, assuming the absence of mutations. One measurement is the expected number of zygotes produced by a type of adult in the ancestral population, multiplied by the survival rate of these zygotes in the descendant population:

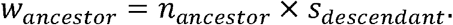

In this case, selection occurs over two generations: fecundity selection in the ancestral population and viability selection in the descendant population. This measurement aligns with the concept of successful offspring frequently used in theoretical biology.

The other way to measure fitness is based on the survival rate of a type of zygote categorized according to genes or traits in the ancestral population, multiplied by the expected number of zygotes produced by that type of adult in the ancestral population:

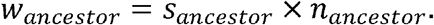

In this case, selection occurs over a single generation, with both viability selection and fecundity selection taking place in the ancestral population. This provides a measurement of fitness in the ancestral population, but it only accounts for the number of zygotes produced by adults and not the number of successful offspring.

## 3 The expansions of the Price Equation

The derivation process of the Price equation can be used to derive a modified partition of the average change in population level, leading to two expansions of the classic Price equation based on the two different measurements of fitness.

### 3.1 Selection happens in two generations

In this scenario, we will examine the process of how adults in the ancestral population generate zygotes that mature into adults in the descendant population. We will assume that there are *N* adults in the ancestral population, labeled with identification numbers *i* = 1, 2,…,*N*, assigned arbitrarily. Let *Z*_*i*_ denote the measurement of interest (such as any trait) for individual *i* in the ancestral population and let 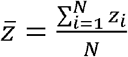 be the average measurement in the ancestral population.

Let *n*_*i*_ be the number of zygotes produced by individual *i*. Then, the average number of zygotes produced by adults in the ancestral population is 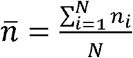, and the total number of zygotes in the descendant population is 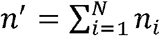. Let 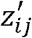 denote the measurement of these zygotes and let 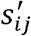 denote the survival rate of these zygotes that mature into adults, with *j* = 1,2,…,*n*_*i*_.

Then, the average measurement of the zygotes produced by individual *i* is 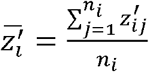. The average measurement of all zygotes is 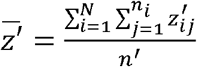. The expected number of adults in the descendant population is 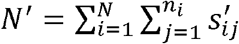, and the average survival rate of these zygotes that mature into adults is 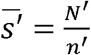. The expected value of the average adult measurement in the descendant population is 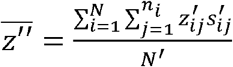. Then, the change of the average measurement after a life cycle is

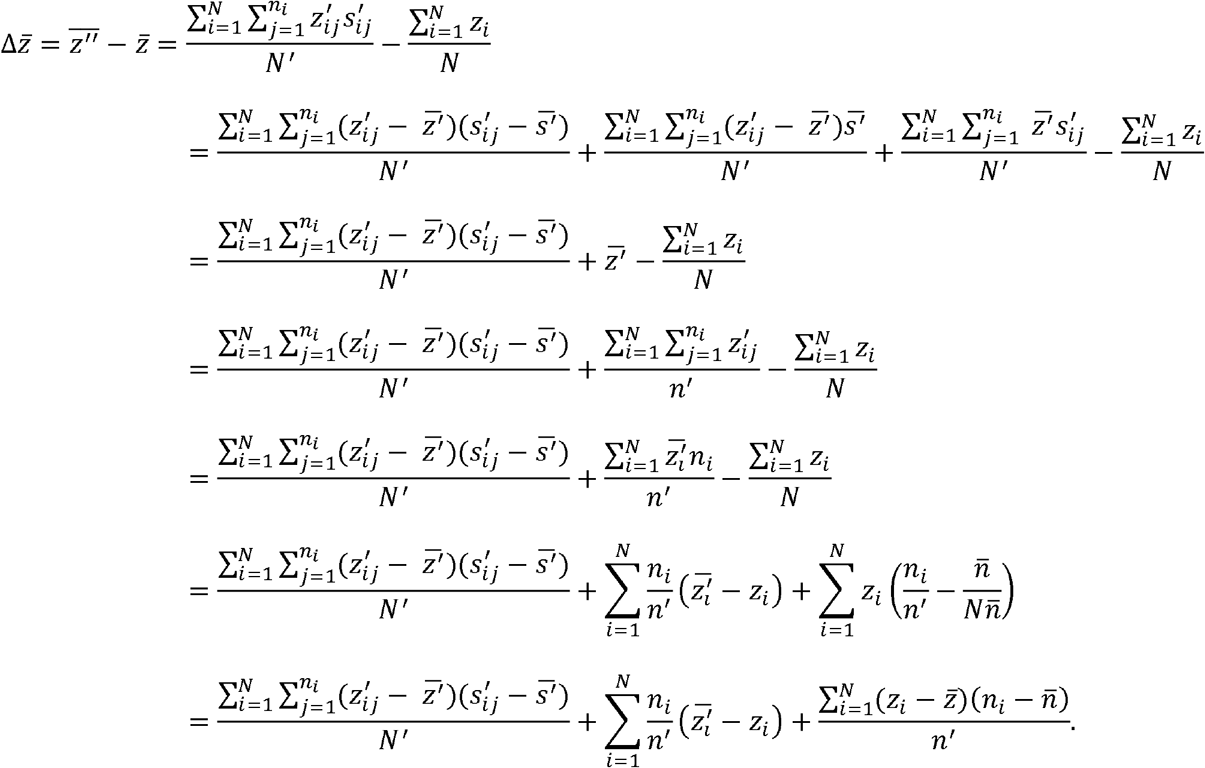

The last line of the equation can be broken down into three terms. The first term, 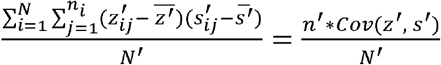, represents the covariance between the measurement of the zygotes in the descendant population and the survival rate of those zygotes, normalized by the expected number of adults in the descendant population. This term captures the effect of viability selection that occurred in the descendant population. The second term, 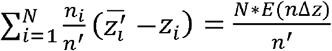 with 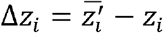, represents the mean of the number of zygotes produced by each adult in the ancestral population multiplied by the difference between the average zygote measurement produced by the adult and the adult’s measurement in the ancestral population, divided by the total number of zygotes in the descendant population. This term captures the effect of mutation, drift, and recombination. Finally, the third term, 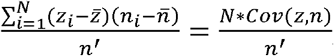 represents the covariance between the adult measurement in the ancestral population and the number of zygotes produced by adults in the same population, normalized by the total number of zygotes in the descendant population. This term captures the effect of natural selection through fecundity selection in the ancestral population.

By switching the order of the terms in the last, we can express the three terms in a more compact form as:

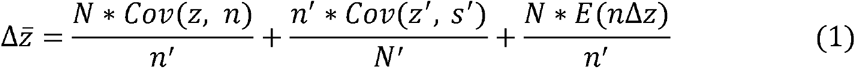

Here, equation (1) shows that the change in the average measurement after a life cycle is influenced by three factors: fecundity selection, viability selection, and mutation, drift, and recombination.

### 3.2 Selection happens in one generation

In this scenario, we examine the process of zygotes maturing into adults in the ancestral population, and these adults producing zygotes that belong to the descendant population. Let the number of zygotes in the ancestral population be denoted as *n*, which are labeled with identification number *i* = 1, 2,…,*n* assigned in any order. Let *z*_*i*_ represent the measurement for zygote *i* in the ancestral population, and the average measurement of zygotes in the ancestral population is 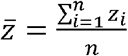. Let *s*_*i*_ be the survival function representing whether zygote *I* matures into a breeding adult successfully or not, such that *s*_*i*_ = 1 if zygote *i* grows up to a breeding adult and *s*_*i*_ = 0 if zygote *i* dies before breeding season. The average survival rate is 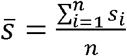. There will be 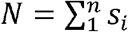 adults in the ancestral population that produce zygotes. Let z^′^_i_ = s_i_z_i_ denote the measurement of adult *i* if zygote *i* could mature into an adult in the ancestral population (*s*_*i*_ = 1). The average measurement of these adults is 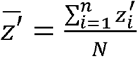. Let *n*_*i*_ represent the number of zygotes produced by individual *i* if zygote *i* could mature into an adult in the ancestral population (*s*_*i*_ =1). Then, the number of zygotes in the descendant population is 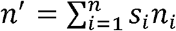, and the average number of zygotes produced by adults in the ancestral population is 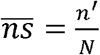. Let 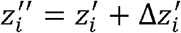, where 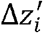 is the difference between each adult’s measurement in the ancestral population and the average zygote measurement produced by the adult, represent the average measurement of zygotes produced by adult *i* in the ancestral population if zygote *i* could mature into an adult in the ancestral population (*s*_*i*_ = 1). The average measurement of zygotes in the descendant population is 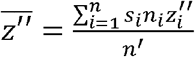. Then, the change in the average measurement after a life cycle is given by:

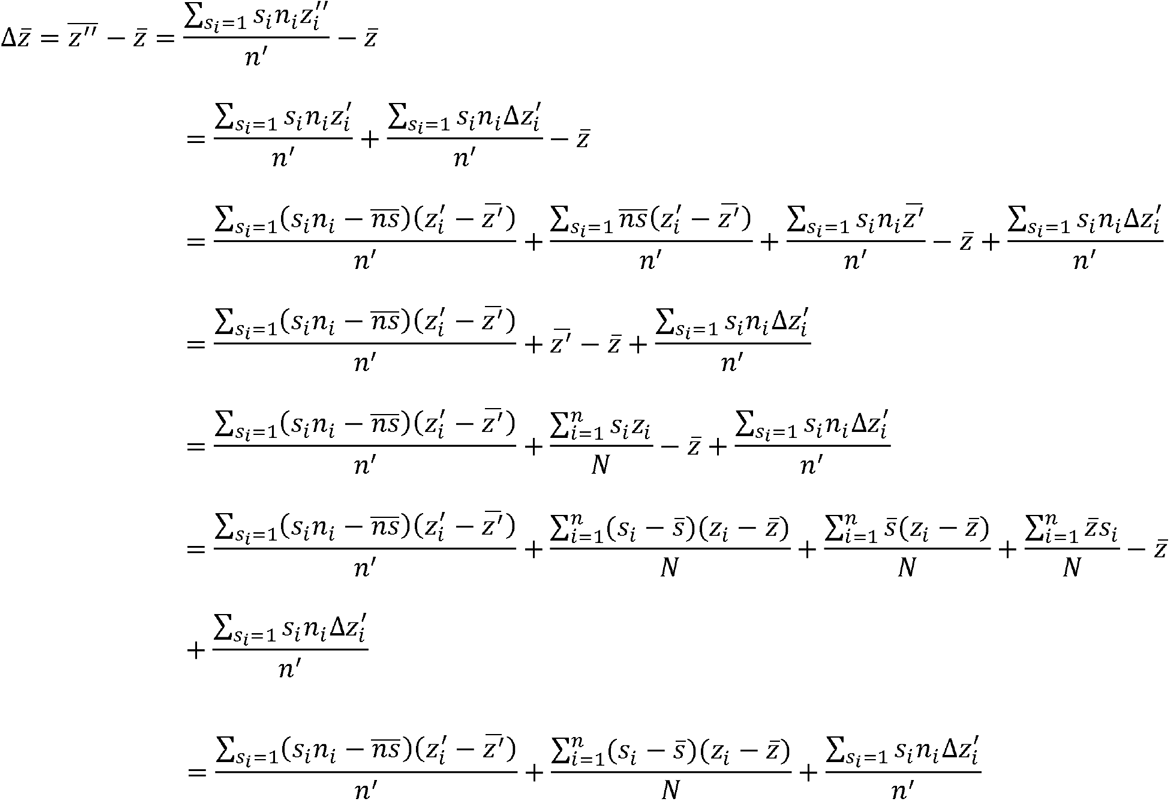

The first term in the last line of the equation, which is 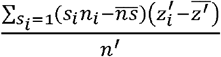, can be expressed as 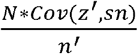 or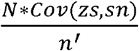. The second term in the last line, which is 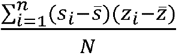, can be expressed as 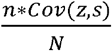. The third term in the last line, which is 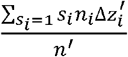, can be expressed as 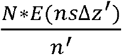, where 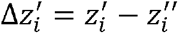. Rearranging the terms in the last line yields equation:

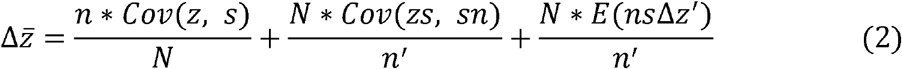

The first term in equation (2) is the covariance between the zygote measurement in ancestral population and the survival function, divided by the total number of zygotes that grow up to adults in the ancestral population. This term captures the effect of viability selection that occurs in the ancestral population. The second term in equation (2) is the covariance between the measurement of adults and the number of zygotes produced by those adults in the ancestral population, divided by the total number of zygotes in the descendant population. This term captures the effect of fecundity selection that occurs in the ancestral population. The third term in equation (2) is the mean of the number of zygotes produced by the surviving adults in the ancestral population times the difference between each adult’s measurement in the ancestral population and the average measurement of the zygotes produced by that adult, divided by the total number of zygotes in the descendant population. This term captures the effect of mutation, drift, and recombination.

### 3.3 Compare with classic Price equation

The classic Price equation has a wide range of applications in theoretical biology, particularly when there is no difference between an ancestor and the average of all its offspring, which implies that the second component of the Price equation is zero. As a result of this assumption, the final terms of equations (1) and (2) are zeros. To compare the quality of the classic Price equation with our expansions, we employ an individual-based model based on the two scenarios used previously to simulate the evolution of life history strategy.

We assume an asexual population with no overlapping generations. Allow every individual to choose a life history strategy for allocating resources for survival and reproduction. We use *x* ∈ [0,1] to symbolize the individual’s strategy of allocating x of its total resources to reproduction and (1-x) of its total resources to survival. The number of zygotes produced is then represented by f(x), while the survival rate from zygote to adult is represented by *g*(1− *x*). We assume that the more resources allocated to reproduction (survival), the more offspring (survival rate from zygote to adult) it has, implying that f(y) and g(y) have positive derivatives. To make things easier, we use *f* (*x*) = *rx*, where r > 0, and *g*(1− *x*)=*s*(1− *x*) ≤ 1, where *s* >0. The survival rate will set to 1 if *g*(− *x*) exceed 1 in the simulation. In this simulation, we additionally suppose that the offspring’s strategy follows normal distribution, with a mean of its parent’s strategy and a variance of *σ*^2^. To ensure that there is no difference between an ancestor and the average of all its descendants, we must ensure throughout the simulation that *rx* ≫ 1. We 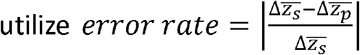, where 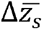 is the average strategy change in simulation and 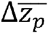 is the predicted average strategy change using the classic Price equation or expansions of the Price equation (Equation (1) or (2)).

When selection occurs in two generations, we simulate an adult population experiencing fecundity selection to create zygotes, and these zygotes experiencing viability selection to grow into adults in the descendant population. The survival adult offspring number is referred to the fitness in the classical Price equation for each individual within ancestral population. The average strategy change in simulation will grow as offspring variety increases. Although the average strategy of zygotes equals the parent’s strategy, the variation of offspring’s strategy may cause the classic Price equation to fail to anticipate the average change in strategy (Figure 1).

**Figure 1.**
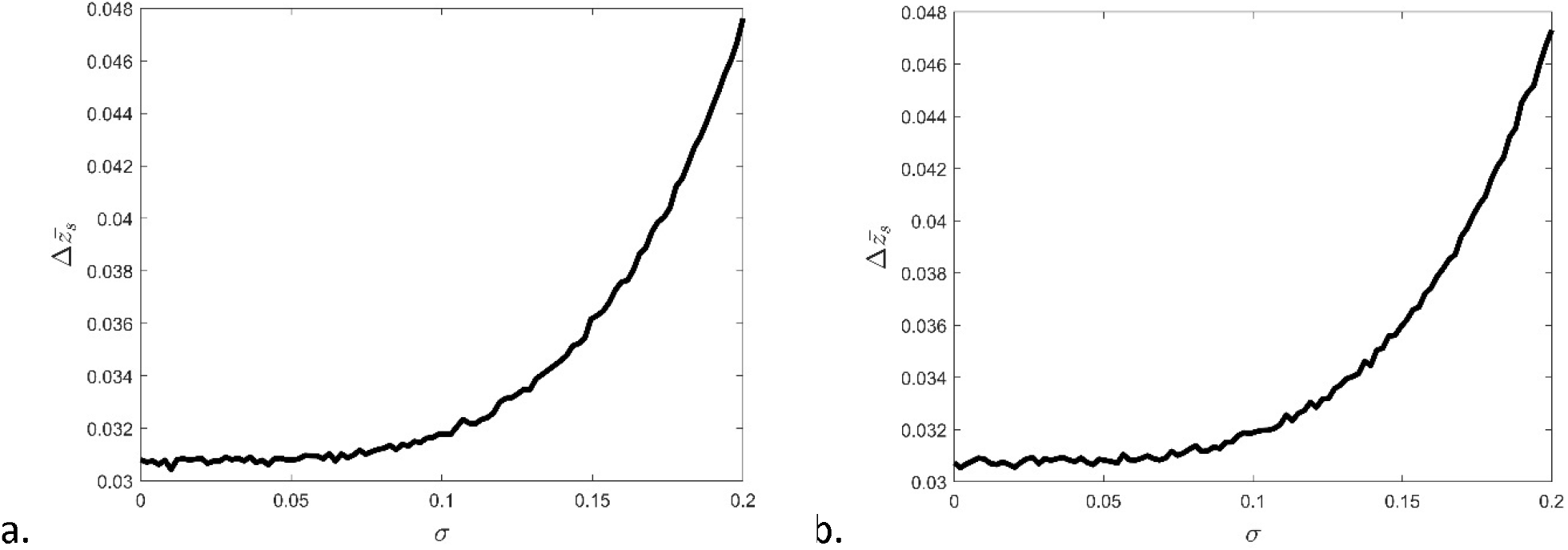
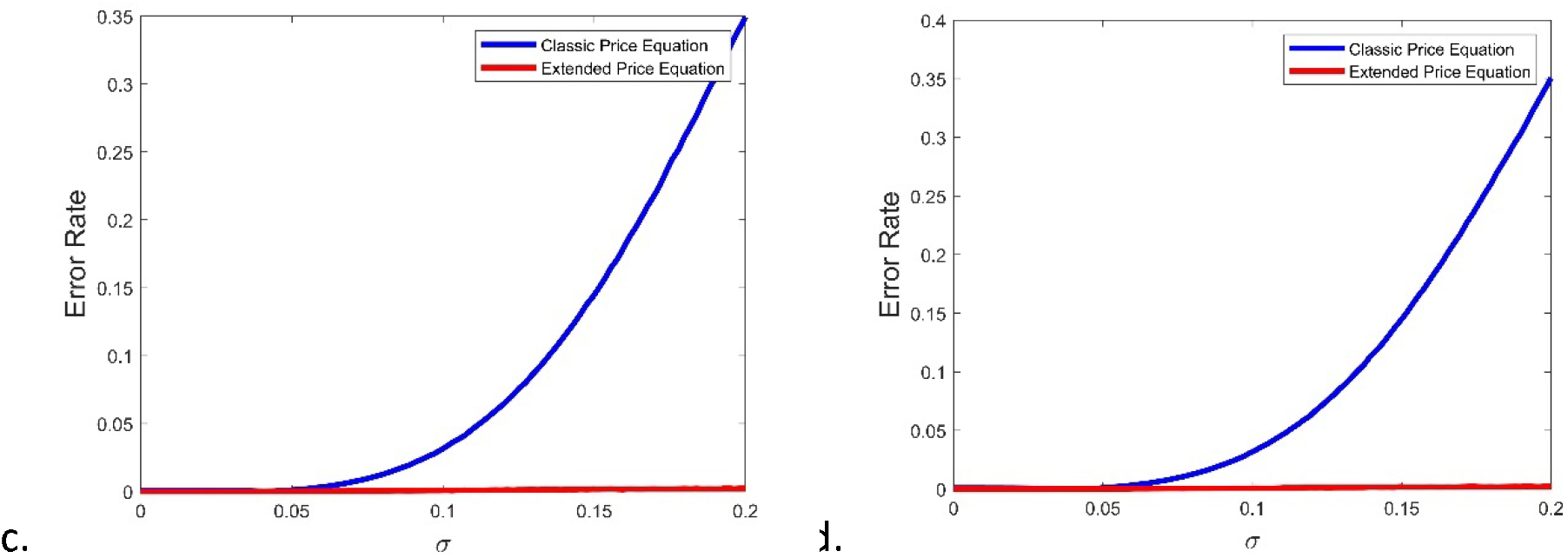
The average strategy changes in simulations and the link between offspring variance and the error rate of the classic and extended Price equations when selection occurs in two generations. The average strategy change in simulation will increase with offspring variation whether **a**. *s*= 0.5 or b. *s*= 5. The error rate in the extended Price equation is near to zero, whereas the error rate in the classic Price equation grows with offspring variation. **c**. *s* = 0.5 and **d**. *s* =5 have no effect on the conclusion. The initial strategy for adults in the ancestry population is chosen at random from [0, 0.2], and we use *r* = 100 to ensure *r* (1 − *x*) ≫ 1. Each simulation is run 100 times, and the average error rate is displayed below.

More importantly, as σ rises, the error rate of the classic Price equation increases. In other words, the classic Price equation couldn’t capture the increasing strategy changes when selection occurs in two generations. However, the expanded Price equation (equation (1)) accurately represents the average change.

The classic Price equation’s weakness in capturing average strategy variation stems from its failure to account for the impact of variation selection on offspring strategies. Notably, although each parent’s strategy equals to the average strategy of its offsprings, offsprings are subjected to variation selection based on their own strategy, which may differ from those of their parents. As a result, offsprings with strategies distinct from their parents face variation selection distinct from their parents, which is not considered in classic Price equation.

When the selection occurs in a single generation, we simulated zygotes being subjected to viability selection and developing into adults, and these adults being subjected to fecundity selection and generating zygotes. The number of zygotes in descendant population is employed to fitness in classic Price equation for each zygote in ancestral population, and it is defined as 0 if the zygote in ancestral population dies before reproduction. In simulation, the average strategy change is unstable. The offspring are not subject to viability selection, and both the classic and extended Price equations can accurately estimate the average change, although the extended Price equation (equation (2)) outperforms the traditional Price equation (Figure 2). All these biases come from the fact that the average strategy of a finite number of randomly selected offspring strategy cannot be exactly equal to the parents’ strategy.

**Figure 2.**
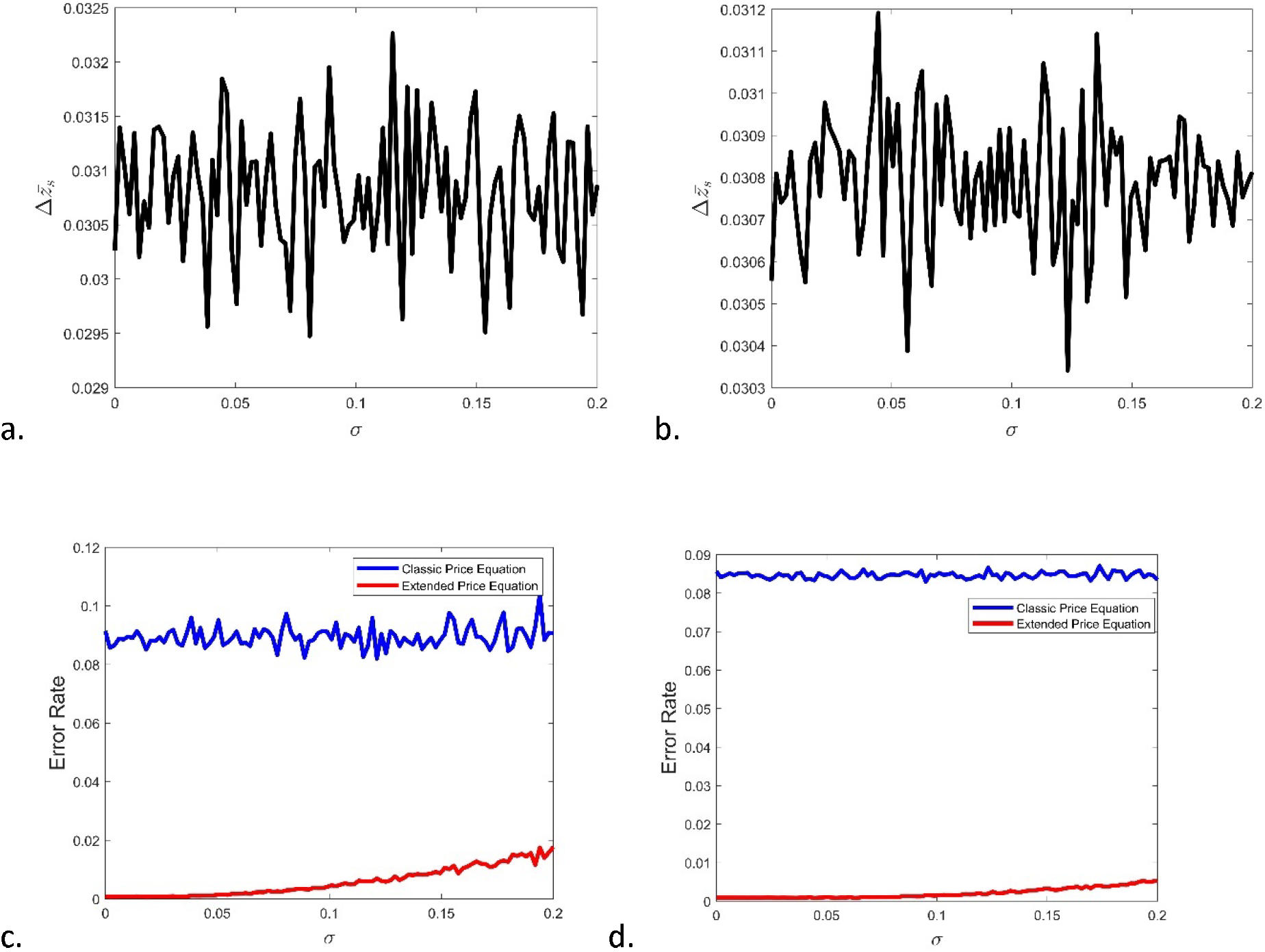
The average strategy changes in simulations and the link between offspring variance and the error rate of the classic and extended Price equations when selection occurs in a single generation. The average strtegy change in simulation fluctuates whether **a**. *s* = 0.5 or **b**. *s* = 5. Both the classic and extended Price equations have error rates near to zero, although the extended Price equation performs somewhat better than the classic Price equation. c. *s*= 0.5 and **d**. *s* = 5 have no effect on the conclusion. The initial strategy for zygotes in the ancestry population is chosen at random from [0, 0.2], and we use *r* = 100 to ensure that *r* (1 − *x*) ≫ 1. Each simulation is run 100 times, and the average error rate is displayed below.

## 4 Discussion

Fitness is a measure of dominance that applies to species, individuals, traits, or genes. Biologists have proposed a multitude of definitions for fitness (Mills & Beatty 1979), which is dependent on the environment but is seldom discussed (Hatfield & Schluter 1999). Different species may undergo distinct forms of selection depending on their environment or at different stages of their life cycle. In this context, we consider a species with a simple life cycle that is subject to viability selection and fecundity selection. Measuring fitness can be challenging, especially when ancestral and descendant populations face varying environmental conditions or selection pressures. When viability selection and fecundity selection occur in two separate generations, fitness is commonly evaluated in terms of the concept of successful offspring in evolutionary biology. This approach, however, is problematic since an individual’s fitness is dependent not just on itself, but also on the survival rate of its offspring. A better way to measure fitness is to consider viability selection and fecundity selection in the same generation, as it provides a more accurate assessment of the population’s fitness based on its own characteristics.

The Price equation offers a way to break down the total change in any biological measurement of interest. While this partition is always accurate, it can be challenging to interpret in terms of selection. Specifically, the mean value term will be zero only if the measurement of an individual and the average measurement of their successful offspring (or zygotes), rather than all zygotes, are equal. Moreover, when a population undergoes both viability and fecundity selection in its life cycle, the classic Price equation’s covariance term fails to capture the distinct effects of each type of selection on the measurement. At times, these selections may even affect the mean value term itself (van Veelen 2020). Nevertheless, the Price equation still offers a useful framework for partitioning.

The classic Price equation focuses on the differences in measurement between the ancestral and descendant populations. However, in our expansions, we consider three populations: the ancestral adult population, descendant zygote population, and descendant adult population (or the ancestral zygote population, ancestral adult population, and descendant zygote population). In the first expansion, where viability and fecundity selection occur in two generations, we use the survival rate to connect the descendant zygote population and descendant adult population and follow the derivation of the classic Price equation to establish the relationship between the ancestral adult population and descendant zygote population. This expansion is a direct expansion of the classic Price equation, as we consider the success of zygotes. In the second expansion, where viability and fecundity selection occur in one generation, we use the survival function to connect the ancestral zygote population and descendant zygote population and follow the derivation of the classic Price equation to establish the relationship between the ancestral adult population and descendant zygote population. In this case, the successful ancestral zygote population is equivalent to the ancestral adult population. These two expansions are essentially the same, as they divide the total change after a life cycle into three parts: viability selection, fecundity selection, and changes in property values between any parent and all its offspring. The third part must be zero if there is no difference between the measurement of an adult individual and the average measurement of its zygotes.

The Price equation has been most useful in the abstract realm, especially in the invariance relations that shed light on the understanding of natural selection. It directly considers fitness or successful offspring/gametes for ancestors, thereby separating out the natural selection term, without distinguishing the roles of viability and fecundity selection. If the first measurement of fitness is used, viability selection in the descendant population can cause differences between ancestral and descendant trait values, rendering the covariance term of the equation inadequate for capturing all the effects of natural selection. Additionally, the assumption of a zero mean value term on the right-hand side of the classic Price equation, commonly made in theoretical work, is invalid (Δ_*z*_ ≠ 0). On the other hand, if the second measurement of fitness is used, it is not appropriate to use successful gametes or offspring. Furthermore, the classic Price equation combines all the roles associated with natural selection and fails to account for important trade-offs between viability selection and fecundity selection that may exist for some traits or genes (e.g., related to resource allocation).

Our expansions are better suited for situations where the environment affects zygotes and adults differently, indicating that viability selection and fecundity selection play distinct roles in the species. For example, when discussing how species allocate limited resources, viability selection and fecundity selection act in opposing ways. The classic Price equation cannot account for these different functions. Kerr & Godfrey-Smith (2009) did provide a more general formulation by expanding the fundamental set mapping between populations. Their expansion can incorporate migration, a feature that our expansions do not have as we focus solely on measuring fitness. Future expansion could explore the integration of these features to make the Price equation more general.

## Acknowledgments

We would like to thank Dolph Schluter for his valuable discussions of this manuscript. We would also like to highlight the comments from Ryo Yamaguchi improved this manuscript a lot. This research was supported by National Natural Science Foundation of China-Yunnan Joint Fund (U2102221).

